# Utilizing virtual experiments to increase understanding of discrepancies involving in vitro-in vivo predictions of hepatic clearance

**DOI:** 10.1101/2020.10.09.332460

**Authors:** Preethi Krishnan, Andrew K. Smith, Glen E.P. Ropella, Lopamudra Dutta, Ryan C. Kennedy, C. Anthony Hunt

## Abstract

We explore a virtual experiment strategy to discover plausible mechanism-based explanations for frequently inaccurate predictions of hepatic drug clearance made using established in vitro-in vivo extrapolation methods. We describe a three-protocol plan that uses validated software analogs of real in vitro and in vivo systems constructed using a common set of objects, components, spaces, and methods. Both systems utilize identical quasi-autonomous hepatocyte analogs containing enzyme-like objects. We parameterize concrete mobile objects (virtual compounds) to simulate the referent drug’s disposition and removal characteristics in vitro and in vivo. The goal of Protocol one (Protocol two) is that measures of virtual compound removal using the in vitro analog (in vivo analog) map directly to the measurements used to compute intrinsic clearance (in vivo hepatic clearance). Protocol three, the focus of this work, requires achieving an essential cross-system validation target. For a subset of virtual compounds, measurements of unbound compounds entering hepatocytes (and their subsequent removal) during virtual in vitro experiments will directly predict corresponding measures made during virtual in vivo experiments. We study four highly permeable virtual compounds when their unbound fraction is fixed at one of seven values (0.05-1.0). Results span the range of hepatic extraction ratios. We achieve the cross-system validation target in 15 cases. In the other 13 cases, explanations of the in vitro-in vivo differences in disposition and removal trace to differences in compound-hepatocyte access within the two analogs during execution. The hepato-mimetic structural organization of hepatocytes within the in vivo analog, which is absent within the in vitro analog, is the determining factor. The results taken together support the feasibility of using the three-protocol plan to help explain observed in vitro-in vivo extrapolation discrepancies. We conjecture that, for some cases, the model mechanism-based explanations of discrepancies described herein will have wet-lab counterparts.

## Introduction

In vitro to in vivo extrapolation (IVIVE) methods, employing hepatocytes or liver microsomes, are widely used in toxicology and during preclinical drug development to predict the hepatic clearance of xenobiotics, particularly in humans. Despite more than a decade of research, reliably accurate predictions are not yet achievable. Underprediction of hepatic clearance is the most daunting problem [1]. Discussions highlight the importance of identifying responsible mechanisms and using that knowledge to develop improved IVIVE methods [1–3]. However, the realities and uncertainties of working with isolated hepatocytes and scaling-derived measures to humans present numerous impediments to disentangling those mechanisms [3,4].

This research’s broadscale objective is to explore the feasibility of using virtual experiments to discover model mechanisms that provide plausible explanations of IVIVE discrepancies. Fig 1 illustrates the plan. It begins with a discrepant IVIVE prediction of hepatic clearance in humans and then follows three protocols. Protocol 1 uses a concretized software analog of an in vitro system employing many virtual hepatocytes. The Protocol’s objective is to obtain a direct quantitative mapping between temporal measures of simulated drug removal and the data used to compute intrinsic clearance.

**Fig 1.**
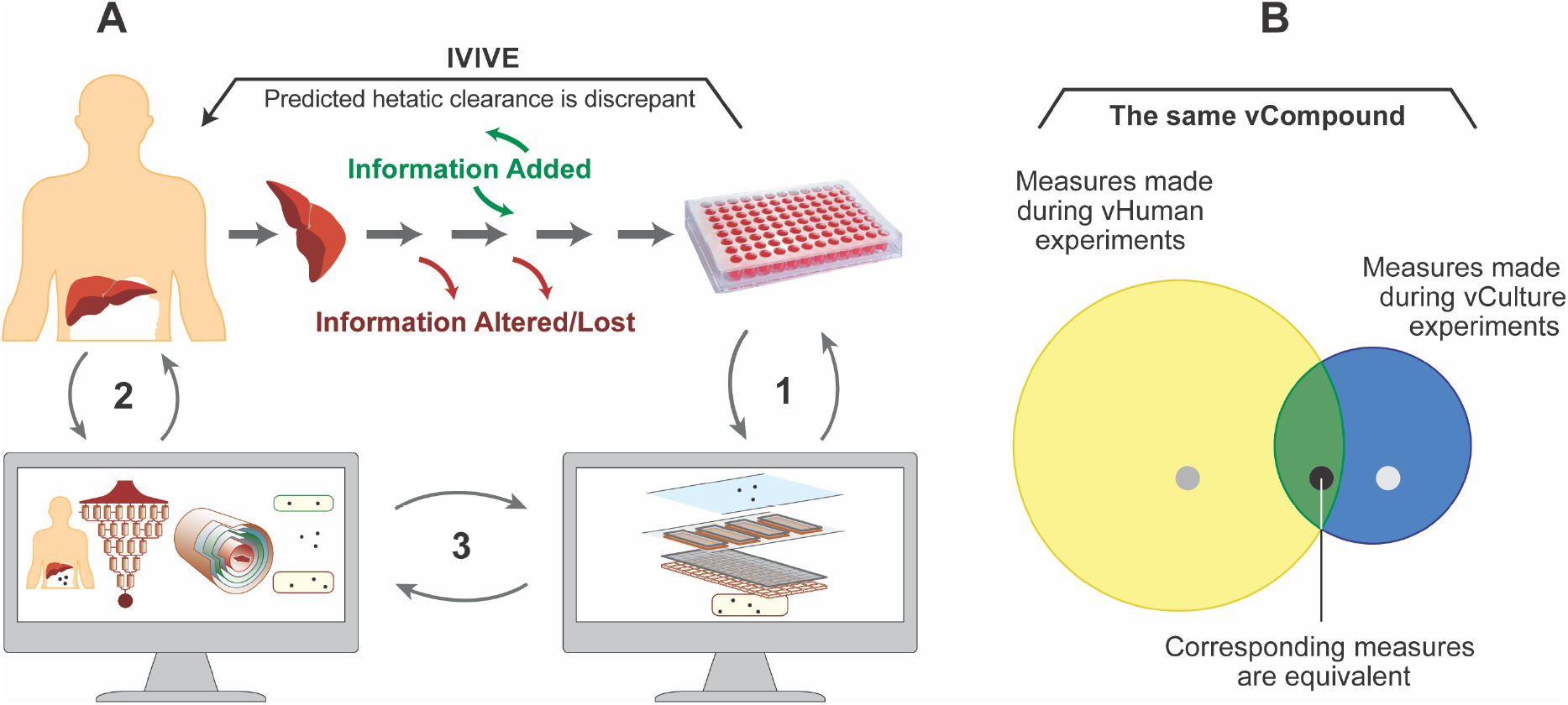
Discovering plausible mechanism-based explanations contributing to IVIVE discrepancies. (A) The plan begins with a failed IVIVE prediction. Protocol 1 establishes a direct quantitative mapping between measures of removal of vCompound during vCulture experiments and the data used to compute intrinsic clearance. vCompound properties mimic selected drug properties. Protocol 2 uses the same vCompound type and is independent of Protocol 1. It establishes a direct quantitative mapping between vCompound removal measures during vHuman experiments and the data used to compute hepatic clearance. Protocol 3 compares and contrasts the details of vCompound disposition and removal during vCulture and vHuman executions. We use the resulting information to posit concrete explanations for the observed IVIVE discrepancy. (B) The blue and yellow circles illustrate the space of possible values for comparable per vHPC measures of vCompound disposition and removal. Measures are quantitatively equivalent within the area of overlap. The black circle illustrates a vCompound that achieves the cross-system validation target (see text), whereas the two grey circles illustrate one that does not.

Protocol 2 uses a concretized software analog of a liver within a human subject. The virtual hepatocytes and simulated drug objects within both systems are identical. The objective for Protocol 2 is to calibrate the simulated drug’s dynamic properties to achieve a direct quantitative mapping between temporal measures of drug removal and the data used to compute hepatic clearance. The purpose of Protocol 3 is to compare, contrast, and explain differences in drug disposition and removal details during Protocol 1 and 2 executions. We envision using the resulting information to posit concrete explanations for the observed IVIVE discrepancy. We design the analogs so that they can be reused without significant structural change, and that is a significant strength of the plan. We can achieve the objectives of Protocols 1 and 2 by adjusting simulated drug properties to reflect those of a target drug [5–9]. Hereafter, to simplify descriptions, we distinguish virtual systems and components from real counterparts by appending the prefix “v,” and we distinguish “virtual” characteristics, properties, and phenomena from real counterparts with capitalization.

We construct vCultures and vHumans independently using a standard set of objects, methods, and modular components. The structure of a vCulture is strongly analogous to that of a typical hepatocyte culture. A vHuman is somewhat analogous to a typical subject used in clinical studies that included measuring hepatic drug clearance. A vHepatocyte (vHPC) object maps directly to actual hepatocytes in vitro and in vivo. Mobile vCompound objects represent the referent drug. The model mechanisms responsible for vCompound disposition and removal during vCulture and vHuman executions are intended to be strongly analogous to their in vitro and in vivo counterparts.

Recent reports document that earlier versions of vCulture and vHuman systems met demanding requirements and explain how they were iteratively refined their model mechanisms so that measurements made during virtual experiments mapped quantitatively to corresponding in vitro and in vivo measurements [8–14]. Having achieved multiple validation targets, the authors argued that the model mechanisms responsible for vCompound disposition/removal and the actual disposition/removal details within their referents were strongly analogous at corresponding levels of granularity. For this work, we build on those arguments using improved versions of both systems.

Consider a particular IVIVE underprediction. Assume that we have completed Protocols 1 and 2, and that direct mappings between temporal measures of vCompound removal and the corresponding wet-lab measures have met prespecified quantitative similarity criteria. At that stage, because we are addressing an IVIVE underprediction, measures of vCompound removal during vCulture experiments will underpredict corresponding measures of vCompound removal during vHuman experiments. The purpose of Protocol 3, which is the focus of this report, is to identify vCulture–vHuman model mechanism differences and determine their causes. Doing so is feasible because we can observe and measure temporal details as they unfold within each system during execution. Together, those differences will support a plausible quantitative mechanism-based theory of explanation for the observed underprediction, one that can be challenged and iteratively improved, as needed. To enable such a theory to become sufficiently credible to influence decision-making during drug development, we must accumulate the requisite supporting evidence. This report provides the initial installment of that evidence.

A core postulate for IVIVE methods is that, under ideal conditions, there is a 1:1 equivalency between in vitro removal rate (clearance) of unbound drug per hepatocyte (or microsomal equivalents) and the in vivo removal rate of unbound drug per hepatocyte. Although it is infeasible to validate that postulate using established in vitro and in vivo methods, it should be feasible to do so for a subset of vCompound properties using vCulture and vHuman experiments. However, we know that changing the structural organization of vHPCs within a vLiver can alter a vCompound’s pharmacokinetic properties, *ceteris paribus* [9]. Hence, an objective for this work that, for a subset of vCompound properties, we confirm a 1:1 equivalency exists between measures of unbound vCompound disposition and removal made during vCulture and vHuman experiments. That 1:1 equivalency serves as a cross-system validation target [15]. Achieving that validation target demonstrates that the area of overlap in Fig 1B exists. In those cases, measures made during vCulture experiments will directly predict corresponding measures made during vHuman experiments. We are also interested in understanding conditions that fail to achieve the 1:1 equivalency, because in those cases, measures made during vCulture experiments will either over- or underpredict corresponding measures made during vHuman experiments. Our working hypothesis is that those failures will be a consequence of temporal differences in model mechanism details that, in turn, will be a combined consequence of vCompound properties and differences in the structural organization of vHPCs within vHuman’s vLiver and within vCulture.

In some cases, those model mechanism-based differences and their explanations may have wet-lab counterparts. We identify vCompound properties that achieve and fail to achieve the cross-system validation target and explain the latter. Taken together, the results support the idea that results of experiments adhering to Protocols 1-3 along with their requirements can help disentangle the mechanisms responsible for IVIVE discrepancies.

## Methods

### The cross-system validation target

Each vCompound type is assigned a unique set of properties (parameterizations, described below). The plan in Fig 1A requires that, for a subset of vCompound types, we accept as valid the core postulate for IVIVE methods mentioned above. Specifically, measures of the per vHPC removal rates of unbound vCompound objects made during independent vCulture and vHuman experiments will be quantitatively equivalent within some similarity criterion, *ceteris paribus*. That equivalency is the cross-system validation target [15] for this work. The area of overlap in Fig 1B illustrates that such equivalencies exist. Each time an unbound vCompound enters a particular vHPC, we record that event. In this work, a vCompound’s fate after it enters a vHPC is independent of the system’s structure and vHPC locations within. Absent a model mechanism that enables a vHPC to “actively” internalize a bound vCompound, vCompound Entry and removal rates per vHPC are directly correlated (when the probability of Exit is constant). Thus, the quantitative equivalency of vCompound Entry rates per vHPC (discussed subsequently) provides alternative evidence that we have achieved the cross-system validation target for that vCompound type.

The area of overlap of the blue and yellow circles in Fig 1B illustrates vCompound types that achieve the cross-system validation target. For those cases, measures of vCompound Entry and removal rates per vHPC will directly predict corresponding measures made during vHuman experiments. The black circle in Fig 1B illustrates an example. The two grey circles illustrate a vCompound type that does not achieve the cross-system validation target. Consequently, measures of its Entry and removal rates per vHPC will either under- or overpredict corresponding vHuman experiment measures.

### Virtual experiments, model mechanisms, and additional requirements

We use previously validated agent-oriented, discrete-event methods. Model execution is a discrete-event Monte Carlo (MC) simulation. The virtual experiment approach involves building, experimenting on, and iteratively refining concrete model mechanisms [6,7,16], while seeking a balance between more detailed biomimicry and the increase in computation programmed into the model systems. Concrete model mechanisms differ fundamentally from the equation-based models [17]. During execution, model mechanisms generate phenomena that we hypothesize can become qualitatively and quantitatively similar to corresponding wet-lab measures when measured. Once we meet the similarity criteria for Protocols 1 and 2, we can claim that vCulture and vHuman model mechanisms are strongly analogous to the real in vitro and in vivo mechanisms. To support such claims, the vCulture and vHuman systems must meet the following requirements.

1. Use absolute grounding [7]. The equation-based models employed by conventional IVIVE methods use absolute grounding, where variables, parameters, inputs, and outputs are in real-world units. Absolute grounding has important advantages and uses, but it limits model reuse and flexibility [5,7,17]. The Fig 1A plan anticipates that we will reuse the vCulture and vHuman systems without significant structural change. Model mechanisms employ relational grounding to deliver the flexibility required and enable model mechanism falsification. Relational grounding requires that variables, parameters, inputs, and outputs are in units defined by other components within each system. Hence, separate quantitative mapping models are required to relate virtual to actual measures. Such model separation increases flexibility and enables the required system reuse. Note that the vCulture–in vitro and vHuman-in vivo quantitative mapping models for different measures will be different necessarily.
2. Components and spaces are concrete and sufficiently biomimetic to facilitate analogical reasoning [18,19].
3. Model mechanisms have context and exhibit the characteristics of an explanatory biological mechanism [20]. Fixed components (e.g., Cells) are arranged spatially and exhibit structure, localization, orientation, connectivity, and compartmentalization. vCompound dynamics mediated by the model mechanisms have temporal aspects, including rate, order, and duration.
4. A vCompound type maps to a particular chemical entity. During each time step, quasi-autonomous components (i.e., software agents such as Sinusoidal Segments (SSs) and vHPCs) recognize different vCompound types and adjust their responses appropriately. For example, a vHPC recognizes that an adjacent vCompound has the property *membraneCrossing* = yes and allows it to Enter (not Enter) stochastically, when other conditions (if any) are met. Parameter names are italicized.

Here, we cite evidence supporting four claims. 1) The parsimonious structural organization of vHPCs within the vLiver (Fig 2) is sufficiently analogous to the histological organization of hepatocytes within human and rodent livers [21,22]. Use-case-specific validation evidence is provided in [12,14,23]. 2) The fine-grain model mechanisms responsible for the removal of vCompounds within vHPCs are concrete, biomimetic, and can be parameterized to be strongly analogous to counterparts in vivo and in vitro. Supportive, use-case-specific evidence is provided in five reports [8,10–12,14]. 3) The structural organization of vHPCs within vCultures is sufficiently biomimetic to simulate the in vitro experiments and measures used to compute intrinsic clearance [8,10,11]. 4) When averaged over many Monte Carlo-sampled executions, mean measures of vCompound dynamics can be scaled to match corresponding wet-lab measurements within prespecified quantitative criteria. Use-case-specific validation evidence is provided in three reports [10,12,14].

**Fig 2.**
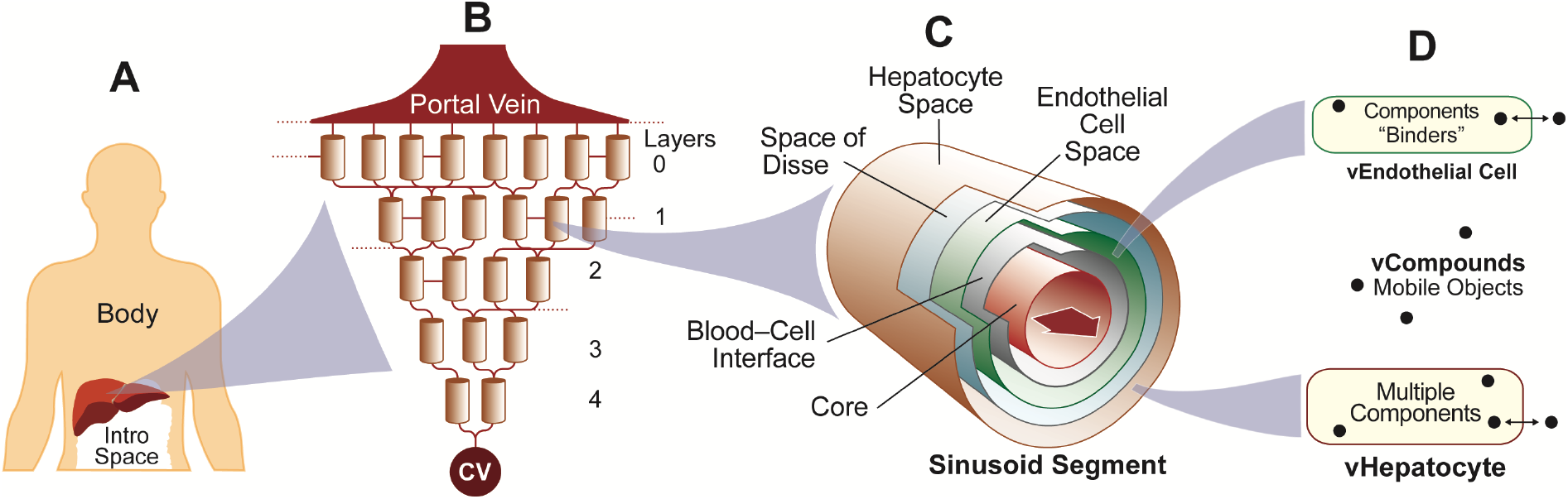
Virtual Human components. (A) A vHuman comprises a well-mixed Body space, a vLiver, and a space for Dose. (B) A portion of vLiver. Graph edges designate flow connections within and between Layers. (C) A multi-layered, quasi-3D Sinusoidal Segment maps to a portion of lobular tissue. It comprises a Core surrounded concentrically by the four 2D grids described in the text. Mobile vCompound objects move within and between these grids. (D) A Sinusoidal Segment contains two Cell types. Each type controls vCompound entry from and exit to an adjacent space. The Cell and the components within determine the fate of vCompounds that enter. We list specification details in Supporting S1 Table.

### Simulation time and vCompounds

During virtual experiments, time advances in discrete time-steps (TS; also called simulation cycles). The duration of each execution is 21,600 TS. For this work, no particular mapping of a TS to real time units is required. In recent work employing virtual acetaminophen hepatotoxicity experiments, the referent organisms are mice [12] and one TS maps to one second. Events occur at a particular instant in time, marking a change of system state. Measurements made at the end of each TS may map to corresponding referent measurements (real or envisioned). However, within a TS, some execution events are codebase dependent and have no direct wet-lab counterparts. We intend that state changes at the conclusion of a TS will be analogous to the net consequences of fine-grain processes that occur in parallel within the referent during the corresponding time interval. To simulate that parallelism, the order of events is randomized for each TS.

We study four vCompound types (vC1-vC4) and Marker. Later, we describe their behaviors during experiments (**vCompound dynamics during experiments**). The Dose for each experiment is 100,000 objects. Marker is always 50% of each Dose, and it serves as a multi-attribute virtual internal standard. Marker does not enter Cells (parameter *membraneCrossing* = false). For vC1-vC4, *membraneCrossing* = true. Mean Marker behavior during repeat executions of the same system is similar, within the variance of MC-sampled executions. As explained in Smith et al., Marker is particularly efficacious during cross-system validation experiments and verification following code changes [12].

vC1-vC4 mimic high permeability xenobiotics. For this work, we had three reasons for limiting attention to only highly permeable vCompounds. 1) We expected that the behaviors of highly permeable vCompounds would make it straightforward to identify vCompound types that achieve the validation target, and 2) make it easier to detect, identify, and correct any inadvertent non-biomimetic differences (having no wet-lab counterparts) between the two systems. 3) By doing so, the duration of executions is reduced. Subsequent work will be needed to establish vHuman-vCulture cross-model verifications for less permeable vCompounds.

### vHuman and vLiver components

Upon initiating an execution, all vHuman and vCulture components are created, assembled and parameterized. We then initiate the experiment protocol and begin measurements. One experiment is a fixed number of MC executions (12 in this work), with a different pseudo-random number seed for each execution. Averaging measures over 12 MC executions is sufficient to detect significant changes in measured phenomena caused by parameter changes. A vHuman (Fig 2A) comprises a vLiver (detailed below), Body, and an Intro space to contain Dose. A vLiver plugs together quasi-autonomous software objects that represent hepatic components at different scales and levels of detail. Microarchitectural features are represented separately from the mechanisms that influence vCompound disposition and Metabolism. A vLiver = 12 MC-sampled vLobules. One vLobule maps to a small random sample of possible lobular flow paths within a whole liver along with all associated hepatic tissue. It is a rough analogy of an actual mammalian lobule, but components are organized to mimic the 3D organization of tissue within actual lobules [8,21]. A directed acyclic graph, with a Sinusoidal Segment (SS) object at each graph node, mediates flow within a vLobule. Flow follows the directed graph edges connecting SSs. Flow paths map to averages of actual flow paths within hepatic sinusoids.

Quasi-3D SS objects are software agents. Each one comprises a Core and five 2D grids arranged concentrically: Blood-Cell Interface Space (simply Interface Space hereafter), Endothelial Cell Space, Space of Disse, Hepatocyte Space, and Bile space (not used in this work and not shown in Fig 2A). An SS in a particular vLobule Layer (described below) functions as an analog of sinusoid components and features at corresponding relative locations averaged across many actual lobules. SS dimensions are MC-sampled, within constraints, at the start of each experiment to mimic lobular variability and simulate a wide variety of Periportal (PP) to Pericentral (PC) flow paths. To minimize differences in vCompound dynamics between vCulture and vHuman experiments, we tightened the constraints on SS dimensions used by Smith et al. [12,14] so that mean SS dimensions in vLivers closely matched the Hepatocyte Space (Fig 3A) dimensions used during vCulture experiments (see **vCulture components**). Thus, SS width is clamped at 15 grid spaces, and the mean length is approximately 5 grid spaces. The mean minimum (maximum) SS length is 3.2 (7.3) grid spaces.

**Fig 3.**
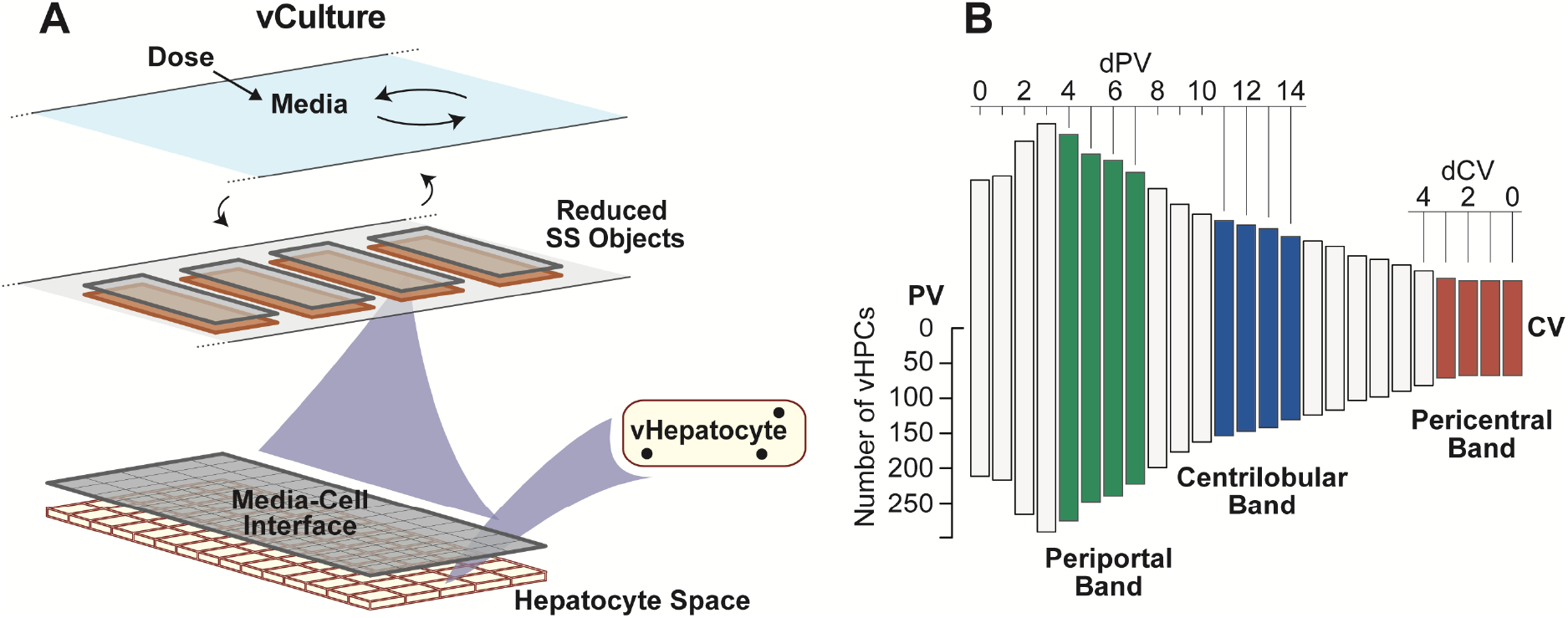
vCulture configuration and relative PV-to-CV vHPC density within vLobules. A vCulture comprises a Media Space plus a single Layer of same-size SS objects (reduced relative to those in Fig 2C). The well-mixed Media Space functions the same as Body. Specification details are listed in Supporting S1 Table. (B) Each bar’s height represents the mean number of vHPCs at the indicated vLobule location averaged over 12 Monte Carlo trials. Moving left-to-right from PV (dPV), the first 15 bars correspond to locations at increasing distances from PV along the average PV-to-CV path. Moving right-to-left, the first 10 bars correspond to locations at increasing distances from CV along the average CV-to-PV path. We average measures within the Periportal, Centrilobular, and Pericentral bands to characterize PV-to-CV differences in vCompound Entry and removal rates per vHPC.

A vLobule has five layers, which can map to lobular zones (Fig 2B). The Portal Vein (PV) connects to all Layer 0 SSs. From the PV to the Central Vein (CV), there are 45, 25, 20, 15, and 9 SS per Layer. There are 55 Layer 0-to-1 edges; 65 Layer 1-to-2; 35 Layer 2-to-3, and 25 Layer 3-to-4 edges. A graph edge connects each Layer 4 SS to the CV. There are also intra-Layer edges (randomly assigned for each execution), which mimic connections between sinusoids: 20 within Layer 0, 7 within Layer 1, 5 within Layer 2, 2 within Layer 3, but none within Layer 4. We fix numbers of intra- and inter-Layer edges for all experiments, but their particular SS-to-SS connections are Monte Carlo-sampled for each execution. Having more edges than SSs enables mimicking the wide variety of PV-to-CV flow paths within lobules [24]. Interconnections are essential to enable the same vLiver to achieve previously described isolated perfused liver validation targets for several different drugs [13,14]. The mean number of vHPCs per vHuman execution = 8,475 (SD = 167). Fig 3B illustrates the mean number of vHPCs at different PV-to-CV locations.

We started with the 3-Layer graph structure used by Smith et al. [12] and increased Layers (from three to five), number of SSs (from 68 to 144), number of graph edges, and reduced mean SS circumferences and lengths by approximately 50%, while maintaining carrying capacity. Those changes improved hepatic disposition biomimicry; specifically, they improved vCompound lateral dispersion and reduced the parallel nature of SS flow paths in Layer 4. The vLobule’s structure was achieved after following the Iterative Refinement Protocol [14,25,26] to achieve qualitative and quantitative cross-validation measures of Marker dynamics and acetaminophen hepatotoxicity measures before and after the above upgrades.

Cell objects are software agents. During each TS, they mediate all interactions with vCompounds, including entry and exit, based on vCompound type properties. Cells occupy 99% Endothelial Cell Space and 100% of Hepatocyte Space. Other Cell object types can be included when required, as in Petersen et al. [10]. Both Endothelial Cells and vHPCs contain binder objects to simulate non-specific binding. They map to a conflation of all cell components responsible for non-specific binding of the referent drug.

In part because we use relational rather than absolute grounding, a vHPC object does not map 1:1 to a hepatocyte (explained in **Component biomimicry has limits**). However, events occurring within vHPCs can map directly to corresponding events believed to occur within hepatocytes, as in Smith et al. [12]. Those events are discussed below in **vCompound Metabolism**.

### vCulture components

The vCulture illustrated in Fig 3A is a partially deconstructed variant of a vHuman. To support achieving the cross-system validation target, we revised the codebase used by Smith et al. to minimize the differences between vCulture and vHuman codebases [12]. The vHuman Intro space and well-mixed Body space are both retained. We rename the latter Media. We reduce vLiver’s directed graph to one Layer (Layer 0 in Fig 2B), which is invariant over MC executions. The single Layer uses 114 Hepatocyte Spaces, which is the same as the number of SSs per vLobule. The Core, Endothelial Cell Space, and the Space of Disse in Fig 2C are not used. We merge the Interface Space, PV, and CV to function as the Media-Cell Interface space (Fig 3A), which is the same size as Hepatocyte Space. Media-Cell Interface can map to the unstirred water layer [27]. One Hepatocyte Space maps to some number of confluent hepatocytes within a hepatocyte culture. For simplicity, all Hepatocyte Spaces are 15 (w) × 5 (l) grid spaces, with one vHPC assigned to each, for total of 8,550 vHPCs, 75 more than the 8,475 in vHuman experiments, a difference of 0.885%.

### vCompound dynamics during experiments

Because vHPC numbers within both systems are the same, essentially, we avoid scaling and make direct graphical comparisons of vCulture and vHuman measures when Dosing in both systems is configured the same, which is the case for all experiments described below. However, when required by Protocol 1 or 2, Dosing can be altered. Upon execution, both systems add Dose to the Intro space. Each TS after that, a fraction of Dose in Intro space is transferred to Body (Media), simulating first order absorption (first order addition to Media). For convenience, we reused the simulated absorption rate used previously for acetaminophen [14].

vCompounds enter the vLiver from Body via the PV. A fraction of vCompounds in Body is moved to the PV each TS to mimic hepatic blood flow. Each TS, vCompounds in the PV are moved randomly to the Core or Interface Spaces in Layer-0 SSs. The parameter *ssFlowRate* controls simulated blood flow in the Core. vCompounds exit an SS via Core and Interface Space and are moved to a lateral or downstream SS along a randomly selected connecting graph edge. Within an SS, extra-Cellular vCompounds percolate stochastically through accessible extra-Cellular spaces influenced by three local flow parameters. Outside the Core, extra-Cellular movement is a biased random walk controlled by the values of *forwardBias* and *lateralBias* listed in Supporting S1 Table. Those values are the same for Marker and vC1-vC4; however, they can be vCompound-specific. vCompounds that exit a Layer-4 SS to the CV are returned to Body. Measurements of vCompound in Body can map quantitatively to measures of a referent drug in plasma (or blood). Because we measure vCompound amounts entering and exiting vLiver each TS, we can compute vLiver Extraction Ratio each TS analogous to how it calculated during perfused liver experiments.

vLiver Extraction Ratio = (vCompounds_PV_ – vCompounds_CV_) / vCompounds_CV_

Each TS during a vCulture experiment, a fraction of vCompounds in Media are transferred to Media-Cell Interface. vCompounds movements within and between Media-Cell Interface and Hepatocyte Space are the same as within and between the Space of Disse and Hepatocyte Space within SSs. vCompounds exit the Media-Cell Interface and return to Media. Because we measure vCompound amounts entering and exiting Media-Cell Interface each TS, we can compute a vCulture Extraction Ratio each TS.

vCulture Extraction Ratio = (amount entering – amount exiting Interface Space) / amount entering Interface Space.

### Focusing on vCompound-vHPC Entry events

vHPCs, their components, and the rules governing component interactions with vCompounds are the same in both systems, but they can be customized as needed during Protocols 1 and 2. In this work, the fate of a vCompound after it enters a vHPC is independent of the system structure and vHPC locations within the system. Each TS, a vCompound that is collocated with a vHPC is allowed to Enter (or not Enter) randomly. An Entry event is a requisite for Metabolism. The value of the vCompound-specific probabilistic parameter *pEnter* determines the occurrence of vCompound-vHPC Entry event (simply Entry event hereafter). Structural differences between vLivers and vCultures can influence the probability each TS of a vCompound being collocated with a vHPC and thus the occurrence of an Entry event.

We record the number of Entry events for each vHPC each TS. Comparing mean Entry rates per vHPC during vCulture and vHuman experiments provides the most reliable method to directly determine the equivalency (or lack thereof) between vCompound disposition and removal dynamics within the two systems. Hereafter, we rely on equivalency of Entry rates per vHPC (hereafter, simply Entry rates) as the cross-system validation target. We also use Entry rate differences to help explain differences in other measures that have wet-lab counterparts, such as the temporal profiles for removal of vCompound from Media and Body.

IVIVE methods typically assume that only the unbound xenobiotic adjacent to a hepatocyte is available to enter. For this work, the probability of a vCompound Entry event (*pEnter*) = (fraction_unbound)×(pEnter_unbound), where pEnter_unbound is the probability of an Entry event for an unbound vCompound. Because the vCompounds in this work represent highly permeable xenobiotics, for simplicity we specify that pEnter_unbound = 1. Thus, hereafter, *pEnter* = fraction_unbound. Each TS, an unbound intra-Cellular vCompound is allowed to exit randomly. The vCompound-specific parameter *pExit* determines the subsequent occurrence of an Exit event.

vCulture-vLiver differences in intra-vHPC mechanisms (Protocols 1 and 2 in Fig 1) and in the ability of the vCompound to Enter vHPCs under comparable dosing conditions may break the 1:1 equivalency and prevent achieving the cross-system validation target. Because vCulture and vLiver intra-vHPC mechanisms are identical in this work, we focus on the degree to which changing *pEnter* may break the 1:1 equivalency.

vC1 and vC2 represent the hypothetical extremes for removing highly permeable xenobiotics. vC1 is not removed; it simply enters and exits all vHPCs. *pExit* for vC1 = 1. vC2 experiences maximal removal; it enters but does not exit vHPCs (*pExit* = 0). vC3 mimics xenobiotics having a small, near zero hepatic extraction ratio. vC4 mimics xenobiotics having a medium hepatic extraction ratio. For vC3 and vC4, *pExit* = 1.

Entry events exhibit no significant location dependency during vCulture experiments. To characterize location-dependent differences in Entry events within a vLiver, we average measures over 12 MC executions within the PP, Centrilobular (CL), and PC bands illustrated in Fig 3B.

Hepatic clearance is defined as the volume of blood that is cleared of drug by the liver per unit of time. By design [7], there is no software counterpart to volume of blood (see **Component biomimicry has limits**). The virtual counterpart to hepatic clearance is the removal rate from Body. Because we employ relational grounding, a separate quantitative mapping is needed to convert the amount of vCompound removed per TS to the volume cleared of an amount of drug per unit of time, as in Petersen et al. [10].

### vCompound Metabolism

Following an Entry event, vC3 or vC4 may be Metabolized. Each vHPC contains Binder objects and four physiomimetic modules [8] that manage events involving Binders: *BindingHandler, MetabolismHandler, InductionHandler*, and *EliminationHandler*. The last two are not used in this work. An Enzyme object is a subtype of Binder objects. An Enzyme can bind and may Metabolize a bound vCompound. In this work, all Binders in vHPCs function as Enzymes. The number of Enzymes in each vHPC is subject to a random draw from *U*(*bindersPerCellMin*, *bindersPerCellMax*), where *bindersPerCellMin* = 5 and *bindersPerCellMax* = 10. Each TS, an unbound vCompound is given an opportunity, determined randomly, to bind to one unoccupied Enzyme. The value of the parameter *pBind* determines whether binding occurs. Upon binding, the vCompound is scheduled to be Metabolized, with probability *pMetabolize*, or released after *bindCycles* = 10 TS. A Metabolized vCompound is deleted and replaced by a Metabolite object. Each TS, a Metabolite is given an opportunity to exit randomly. The subsequent occurrence of an Exit event is determined by its value of *pExit*. *pExit* for all Metabolites = 1; however, when needed, *pExit* can be made Metabolite-specific. After exiting a vHPC, a Metabolite does not reenter Cells. To facilitate direct comparisons of results of experiments using different vCompound types, the parameters controlling Metabolite movement within and between extra-Cellular spaces in both systems are the same as for vC3, vC4, and Marker. When needed, we can employ multiple Enzyme types [12,14]. In those cases, *pBind*, *bindCycles*, *pMetabolize*, and *U*(*bindersPerCellMin*, *bindersPerCellMax*) can be Enzyme-type and vCompound-type-specific.

### Making virtual measurements analogous to wet-lab measurements

We measure virtual features and phenomena analogous to how corresponding wet-lab measurements are (or might be) made. Doing so strengthens the virtual-to-wet-lab experiment analogy. Many parameters are probabilistic. Specifications of several features are MC-sampled at the start of each execution. Because we average measurements over 12 MC executions, some may exhibit considerable variability. That variability is intentional. It represents and helps account for the variability and uncertainty that characterizes wet-lab measurements.

### Component biomimicry has limits

None of the objects in Fig 2 and Fig 3A are intended to model actual biological counterparts explicitly. Instead, their organization, function, and behaviors during execution—the model mechanisms—are intended to be sufficiently analogous to their biological counterparts so that prespecified fine- and coarse-grain measures, recorded during executions can map quantitatively to corresponding Protocol 1 and 2 validation targets [17].

An SS does not map directly to a portion of a single sinusoid and adjacent tissue. Instead, events occurring within a particular SS are intended to be strongly analogous to corresponding events occurring at corresponding relative PV-to-CV locations. The mapping from cylindrical 2D Hepatocyte Space in Fig 2C to corresponding 3D configurations of hepatocytes is an approximation. A vHPC at a particular PV-to-CV location within a vLiver maps to a random sample of hepatocytes (or hepatocyte functionality) accessed by referent drug at a corresponding relative PV-to-CV location. Each vHPC within a vCulture maps to a same-size random sample of hepatocytes (or hepatocyte functionality), ideally isolated from a referent liver. Thus, a vHPC cannot map 1:1 to a hepatocyte, although there are strong functional analogies. It follows that a vLobule does not directly model liver microanatomy, yet its contribution to model mechanisms is intended to be hepato-mimetic during execution.

### Hardware and software details

The Java-based MASON multi-agent toolkit was used to develop the vLiver and vCulture. Experiments were executed using local hardware running 64-bit Linux Mint and Google compute engine was used as the virtual machine. The virtual framework was created using Java. The R programming language was used for analyses and plotting data. vHumans, vCultures, and configuration files are managed using the Subversion version control tool in two repositories, one private (Assembla) and another public. Values for key vHPC specifications and parameterizations are listed in Supplemental Table S1. Quality assurance and control details, along with practices followed for validation, verification, sensitivity analyses, and uncertainty quantification, areas discussed in Smith et al. [14]. The toolchain, operating system, configurations, and our entire codebase is available on (https://simtk.org/projects/isl/).

## Results

All results from experiments using vC1 (first subsection) achieve the cross-system validation target. Results from experiments using vC2 achieve the cross-system validation target for *pEnter* = 1 (second subsection) but fail to do so for *pEnter* ≤ 0.8. We provide model mechanism-based explanations for those failures. vC3 (third subsection) achieves the cross-system validation target for all values of *pEnter*. In the final subsection, we describe how and why vC4 fails to achieve the cross-system validation target for all values of *pEnter*.

### Achieving the cross-system validation target using vCompound-1

Figures 4 and 5 contain mean measures of vC1 during vCulture and vHuman experiments. Under dynamic steady-state conditions, for *pEnter* = 1.0, the mean plateau values for percent of Dose in Media (Fig 4A) and vHuman Body (Fig 5A) are equivalent within the variance of 12 MC executions. The pattern of changing plateau values is similar when using smaller *pEnter* values. However, for larger *pEnter* values, the vCulture plateau values are smaller than corresponding values from vHuman experiments.

**Fig 4.**
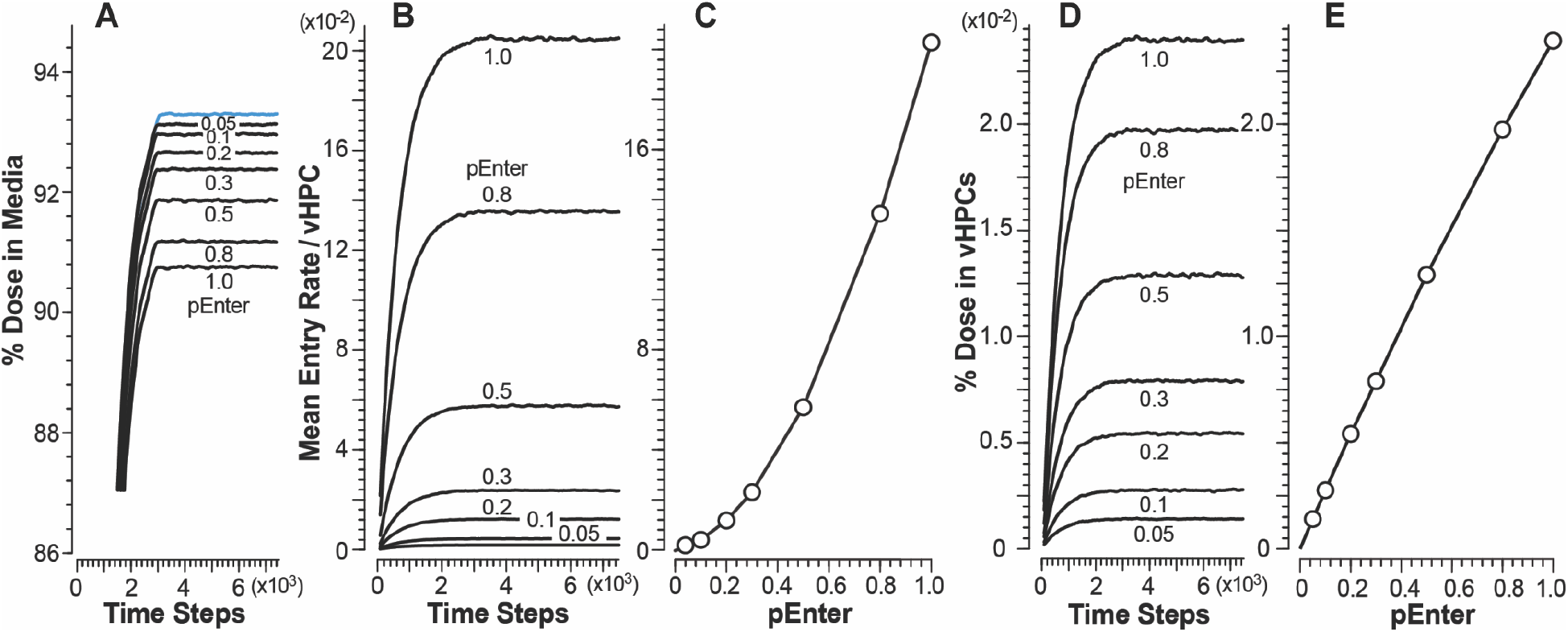
Results from vCulture experiments using vC1. Changing *pEnter* alters vCompound dynamics during each experiment. Temporal values here, and in the subsequent figures, are centered moving averages spanning 181 TS. (A) Temporal measures of percent Dose in Media for each *pEnter*; blue measures are for Marker. (B) Temporal measures of mean Entry rates. (C) Mean dynamic steady-state Entry rates for each *pEnter*. (D) Temporal measures of percent Dose and (E) mean dynamic steady-state values of percent Dose within all vHPCs for each *pEnter*.

**Fig 5.**
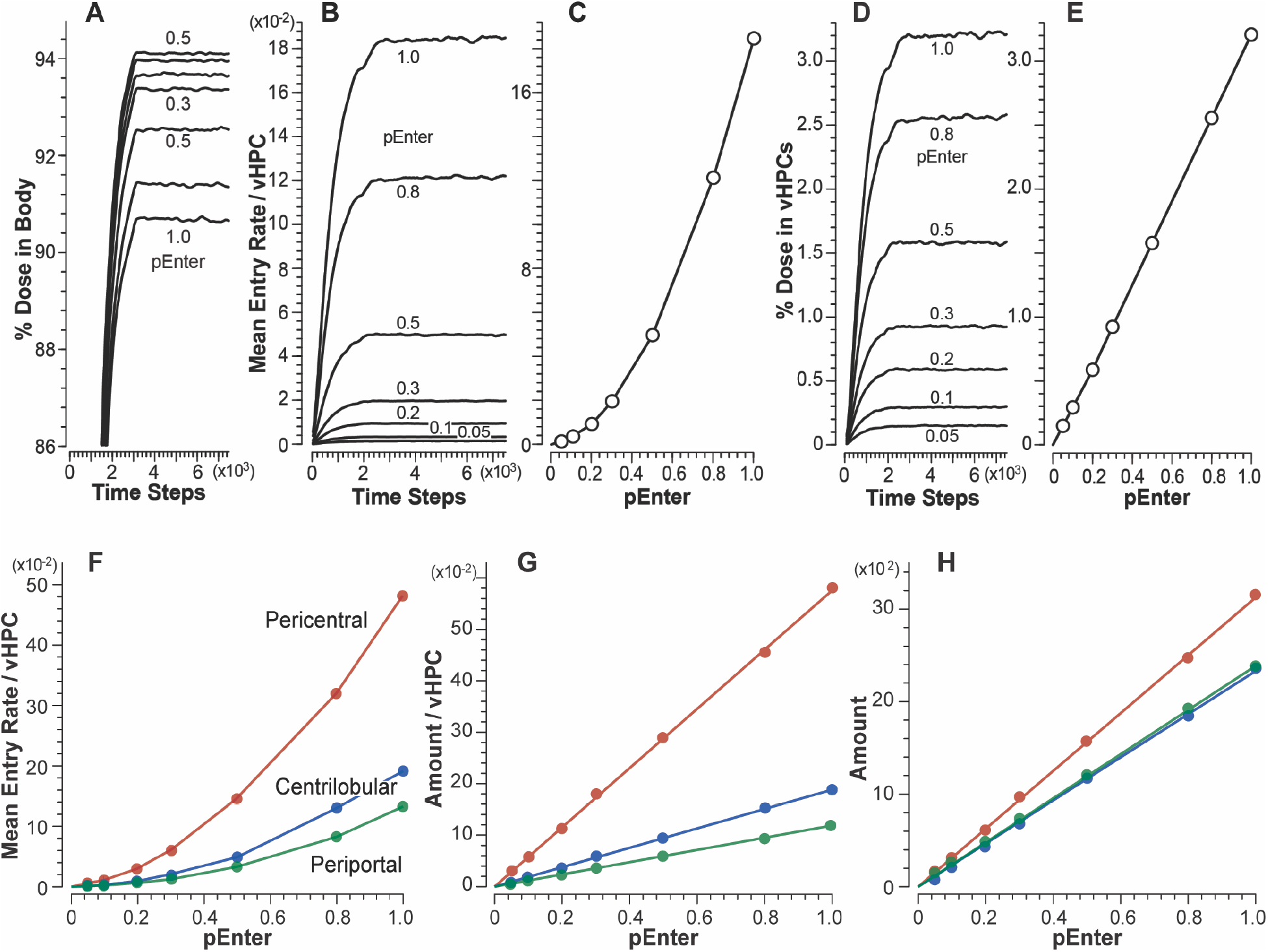
Results from vHuman experiments using vC1. (A) Temporal measures of percent Dose in Body for each *pEnter.* (B) Temporal measures of mean Entry rates for each *pEnter*. (C) Correlation between mean dynamic steady-state Entry rates and *pEnter*. (D) Temporal measures of percent Dose in vHPCs for each *pEnter.* (E) Correlation between mean dynamic steady-state amounts of vC1 in vHPCs and *pEnter*. Measures in F-H are mean dynamic steady-state values within the Periportal, Centrilobular, and Pericentral bands in Fig 3B. (F) Correlation between mean dynamic steady-state Entry rates and *pEnter*. (G) Correlation between vC1 amounts per vHPC and (H) total amounts within all vHPC and *pEnter*.

The mean plateau Entry rates in Fig 4B and Fig 5B decrease with decreasing values of *pEnter*. For each *pEnter*, the mean Entry rates are smaller in vLiver (most evident for *pEnter* = 1.0) because a small fraction of vC1 within Core, Interface Space, Endothelial Cell Space, and the Space of Disse exits to CV without accessing Hepatocyte Space. Hence, the accessibility of those additional spaces reduces slightly the dynamic exposure of vCompounds to vLiver vHPCs relative to vCulture vHPCs. After considering exposure difference and the variance across 12 MC executions, mean plateau vC1 Entry rates for all *pEnter* values are equivalent in both systems. Thus, we achieve the cross-system validation target, and vCulture Entry rates adequately predict corresponding vHuman Entry rates for all *pEnter.*

The correlations between Entry rates and *pEnter* (Fig 4C and Fig 5C) exhibit the same nonlinearity. After entering vCulture’s Media-Cell Interface Space (or the Space of Disse within a vLiver SS), a vC1 using *pEnter* = 0.5-1.0 may enter and exit more than one vHPC before exiting to Media (or exiting that SS). The probability of multiple Entry events is reduced considerably for vC1s using smaller *pEnter* values. Under dynamic steady-state conditions, we expect vC1 amount per vHPC to be directly proportional to *pEnter*. Correlations between amounts in vHPCs and *pEnter* (Fig 4E and Fig 5E) verify that expectation.

Mean plateau vC1 Entry rates and amounts per vHPC are the same for all vCulture vHPCs. That is not the case for vHuman experiments (Fig 5G-H), where measures depend on the vHPC’s PV-to-CV location. Because the number of vHPCs decreases PV-to-CV (Fig 3B), the correlation between mean Entry rates and *pEnter* (Fig 5F) and between mean amounts per vHPC and *pEnter* (Fig 5G) is largest within the PC band and smallest within the PP band. For *pEnter* = 1.0, mean plateau Entry rates increase 1.4-fold. (3.6-fold) from the PP to the CL (PC) band. The increase is larger for smaller *pEnter* values. For example, when using *pEnter* = 0.1, mean Entry rates increased 1.7-fold (5-fold) from the PP to the CL (PC) band. The explanation for that nonlinear pattern is the same as that provided above for Fig 4C and Fig 5C. Although the amount per vHPC is larger within the CL band than within the PP band, there are fewer vHPC within the CL band. Consequently, mean amounts within the two bands are similar (Fig 5H). In similar experiments that otherwise have a constant PV-to-CV vHPC density (analogous to the conventional parallel tube liver model), mean Entry rates and amount of vC1 per vHPC would decrease PP-to-CL-to-PC.

### Achieving and failing to achieve the cross-system validation target using vCompound-2

Fig 6 and Fig 7 show measures of vC2 disposition and removal during vCulture and vHuman experiments. For vC2, because *pExit* = 0, an Entry event is also a removal event. After taking into account that Interface Space, Endothelial Cell Space, and the Space of Disse are absent in vCultures, the temporal profiles (and areas under each curve) for percent dose in Media (Fig 6A) and Body (Fig 7A) are equivalent for *pEnter* = 1.0 within the variance of 12 MC executions. The Entry rate profiles (Fig 6B and Fig 7B) for *pEnter* = 1.0 are also equivalent. Thus, for *pEnter* = 1.0, the cross-system validation target is achieved: vCulture measures predict corresponding vHuman values. However, for *pEnter* = ≤ 0.8, the cross-system validation target is not achieved: vCulture measures underpredict corresponding vHuman measures and the magnitude of the underprediction increases as *pEnter* decreases.

**Fig 6.**
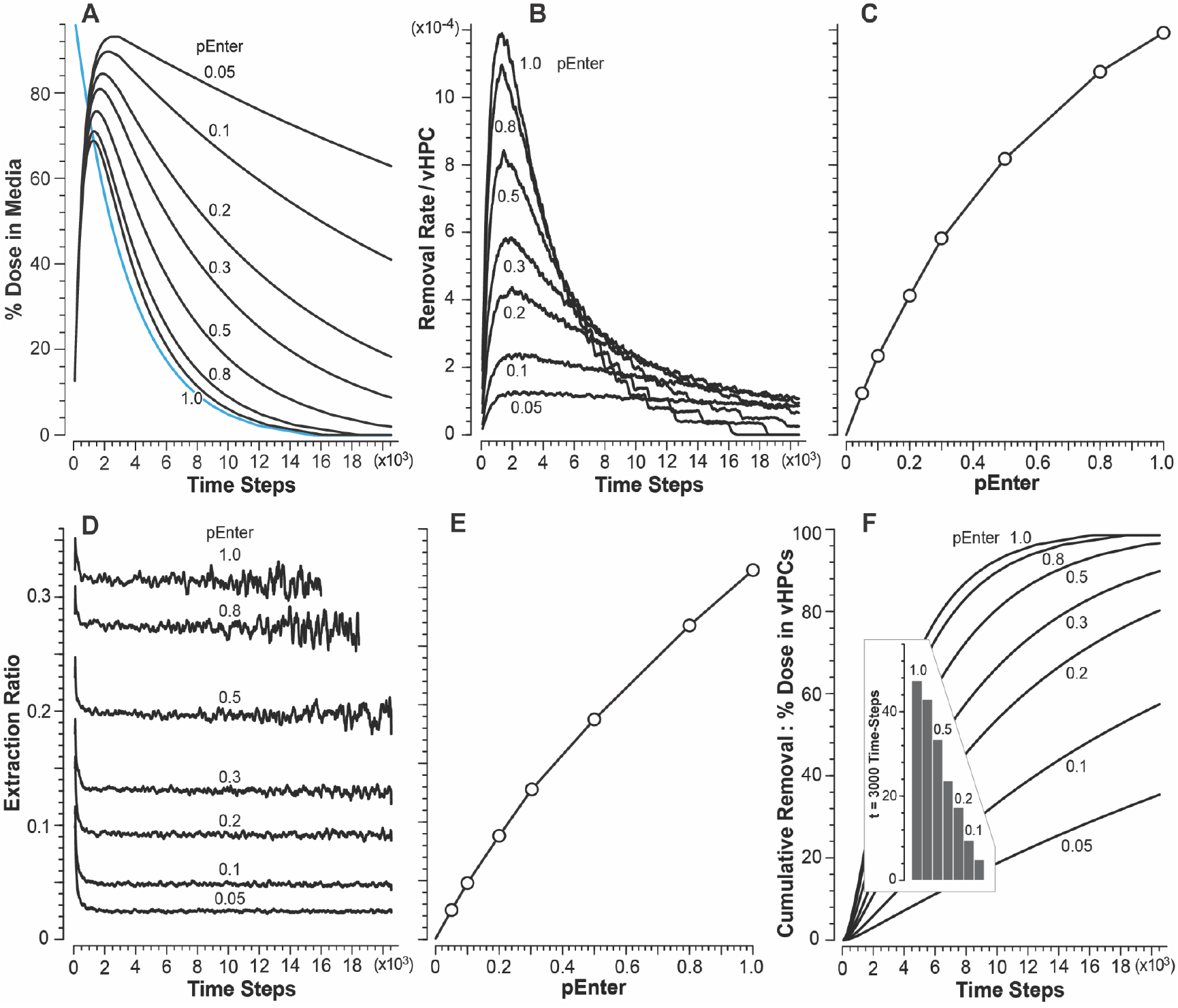
Results from vCulture experiments using vC2. (A) Temporal measures of percent Dose in Media for each *pEnter.* Blue profile: all vC2 added to Media at t = 0. (B) Mean removal rates for each *pEnter* (Entry rates = removal rates). (C) Correlation between mean peak Entry rates and *pEnter*. (D) Temporal measures of Extraction Ratio for each *pEnter*. The variance increases with time because smaller amounts are measured each TS. Extraction Ratios for *pEnter* = 1.0 and 0.8 terminate because (essentially) all vC2 has been removed. (E) Correlation between mean plateau Extraction Ratios and *pEnter*. (F) Temporal measures of percent Dose in vHPCs for each *pEnter*. Insert: mean values at t = 3,000 TS for each *pEnter*.

**Fig 7.**
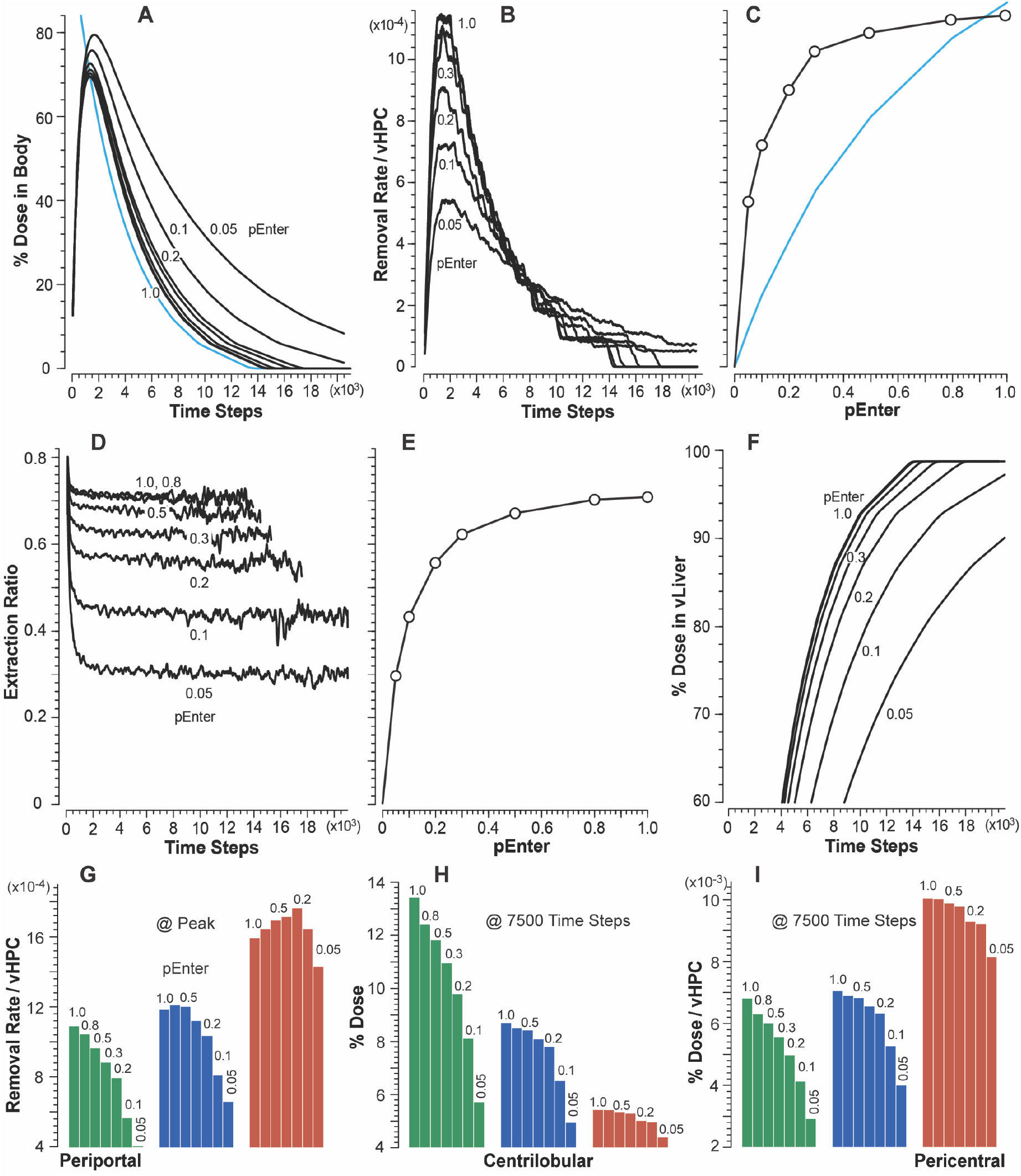
Results from vHuman experiments using vC2. (A) Temporal measures of percent Dose in Body for each *pEnter.* Blue profile: all vC2 was added to Media at t = 0. (B) Mean Entry rates for each *pEnter* (Entry rates = removal rates). (C) Correlation between mean peak Entry rates and *pEnter*. To facilitate comparisons, the blue curve traces the corresponding values from Fig 6C. (D) Temporal measures of Extraction Ratio for each *pEnter.* For p ≥ 0.2, measures terminate early because (essentially) all vC2 has been removed. The variance increases with time because smaller amounts are measured. (E) Correlation between mean plateau Extraction Ratios and *pEnter*. (F) Temporal measures of percent Dose in vLiver for each *pEnter*. (G) Peak Entry rates for each *pEnter* within the three bands in Fig 3A. Corresponding measures of percent Dose (H) and percent Dose per vHPC (I) at 7500 TS.

During vCulture experiments, the correlation between mean peak vC2 Entry rates (averaged over 1,000 TS) and *pEnter* (Fig 6C) is concave. Conventional theory is that it should be linear. The Entry rates for *pEnter* ≤ 0.8 are larger than one might expect. The reason for the nonlinearity: when a vC2 (using *pEnter* ≤ 0.8) adjacent to a vHPC in Hepatocyte Space is not allowed to enter, when given an opportunity, it may have one or more additional opportunities to enter a vHPC before returning to Media. The net consequence of those stochastic events is analogous to the unstirred water layer effect described by Wood et al. [27]. The nonlinearity in Fig 6C is also evident in percent Dose remaining in Media (Fig 6A), Extraction Ratios (Fig 6D and Fig 6E), and cumulative removal (Fig 6F).

When using *pEnter* ≤ 0.8, measures of percent Dose in Media (Fig 6A) and cumulative removal (Fig 6F) during vCulture experiments underpredict corresponding measures during vHuman experiments. It is noteworthy that measures in Fig 7A-D and Fig 7F are relatively robust to changes in *pEnter* within the 0.5-1.0 range. For vCulture experiments using *pEnter* = ≤ 0.5, mean peak Entry rates (and areas under percent Dose in Media curves) considerably underpredict corresponding vHuman measures (Fig 7B). An example for *pEnter* = 0.3 (0.1; 0.05), the mean peak Entry rate is underpredicted by 1.4-fold (3.1- and 4.4-fold). Those underpredictions are a consequence of the tapered structural organization of vHPCs within vLivers (Fig 2B) being absent in vCultures. The fine-grain events occurring within PP, CL, and PC bands explain how and why measures made during vCulture experiments underpredict corresponding vHuman measures, even though vHPCs are identical in both systems.

The dependency of mean vC2 removal rates on location within vLivers (Fig 7G-I) is a consequence of upstream removal of vC2 (Fig 7H), the fact that the number of vHPCs decreases PV-to-CV, and the intra-Layer edges connecting SSs within Layers 0-3. The latter causes the length of the PV-to-PC path taken by some vCompounds to be much longer than the shortest PV-to-PC path. The magnitude of the differences within each of the three bands increases with decreasing *pEnter*. To illustrate, for *pEnter* = 1.0 (0.05), the PC/PP ratio for peak Entry rates (Fig 7G) is 1.4 (3.5). For percent of Dose per vHPC at *t* = 7,500 TS (Fig 7I) when using *pEnter* = 1.0 (0.05), the PC/PP ratio is 1.5 (2.8). The consequences of changing *pEnter* on vHPC Entry rates and cumulative removal are clearly evident within the PP band (Fig 7G,H). The influence of decreasing *pEnter* diminishes within the CL band and is almost absent within the PC band. Although the amounts of vC2 removed within the three bands decrease PP-to-PC, the per vHPC removal rates Fig (7G), and thus per vHPC amounts (Fig 7I), increase PP-to-PC, which means that the PC vHPCs do more of the vC2 removal work, unlike in vCultures. Those observations may aid in disentangling explanations of IVIVE underpredictions.

In both vCulture and vHuman experiments, a step-like pattern is evident in post-peak measures of removal rate (Fig 6B and Fig 7B). That pattern is a consequence of specifying that the mean SS length in vHuman experiments be approximately 5 grid spaces, which is the same as the length of all Hepatocyte Spaces used by vCulture. Making those lengths equivalent results in both systems containing similar numbers of vHPCs, facilitating direct comparisons of results from vCulture and vHuman experiments. When each SS’s length is MC-sampled from a wider distribution, that step-like pattern vanishes [14].

### vCompound-3 achieves the cross-system validation target

Measures of vC3 disposition and Metabolism during vCulture and vHuman experiments are provided in Fig 8. vC3 represents a highly permeable but very slowly metabolized xenobiotic. It uses *pBind* =0.01 and *pMetabolize* = 0.005. Otherwise, the parameters that determine vC3 dynamics in both systems are identical to those used by vC1. For all *pEnter*, the percent of Dose in Media and Body (Fig 8A) are equivalent within the variance of 12 MC executions. Metabolite accumulates faster in Media than in Body (Fig 8B) because Metabolite in vLiver must traverse additional spaces and downstream SSs before exiting to CV and moving to Body.

**Fig 8.**
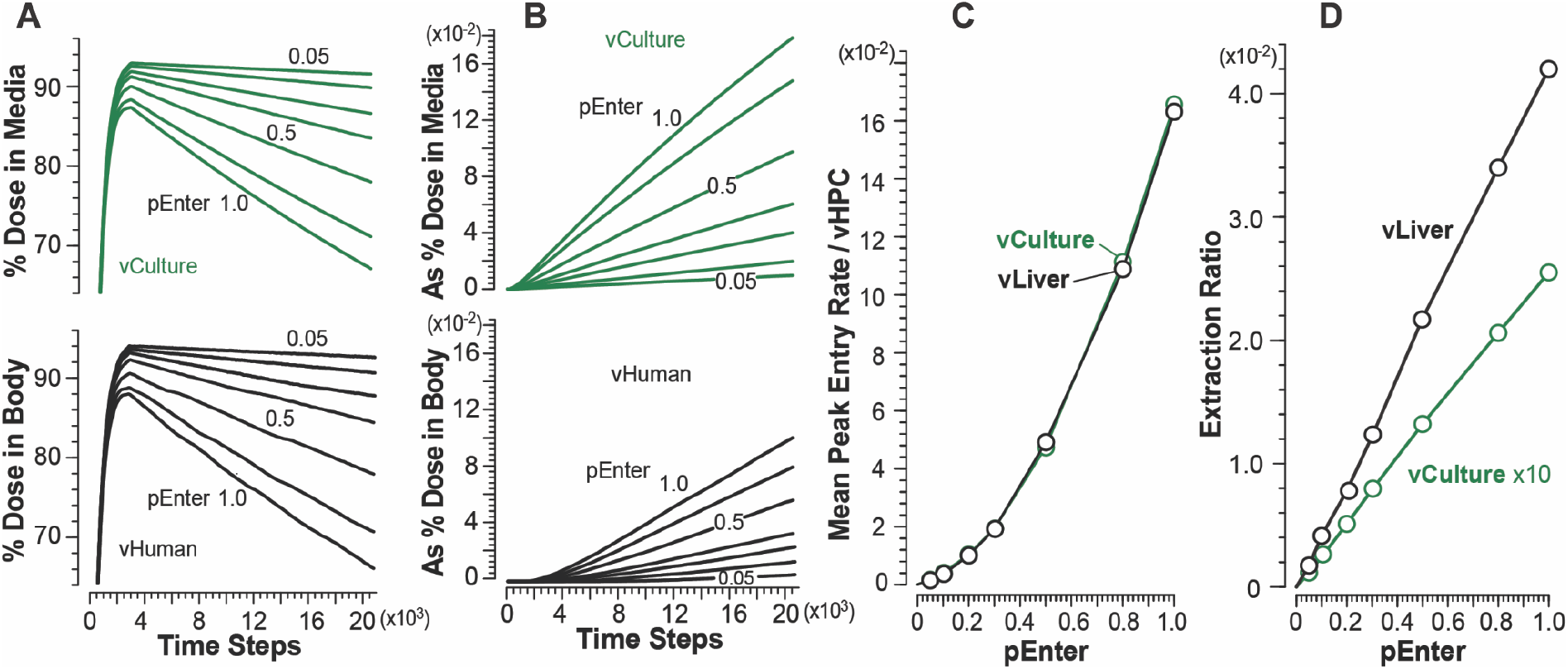
Results from vCulture and vHuman experiments using vC3. (A) Temporal measures of percent Dose in Body (top) and Media (bottom) for each *pEnter.* (B) Amount of Metabolite in Body (top) and Media (bottom) as percent of Dose for each *pEnter.* (C) Correlations between mean peak Entry rates and *pEnter*. (D) Correlations between mean Extraction Ratios and *pEnter*.

We can infer from vC1 results that, because vC3’s rate of Metabolism is small, Entry rates will be approximately equivalent in both systems, which is the case (Fig 8C). The explanation for the nonlinearity in Fig 8C is the same as that provided above for vC1. Thus, the cross-system validation target is achieved for all *pEnter*, and vCulture measures directly predict corresponding vHuman measures.

During vHuman experiments, mean peak Entry rates increase PV-to-CV similar to increases measured for vC1 (Fig 5F). To illustrate for *pEnter* = 1.0 (0.1), the PC/PP ratio for mean peak Entry rate is 3.6 (4.4). The expected large between-system difference in mean Extraction Ratios (Fig 8D) is a combined consequence of three features. 1) The upstream Binding of vC3 to Enzymes. 2) The number of vHPCs decreases PV-to-CV, which increases downstream per vHPC exposures, as demonstrated in Fig 5F-H. 3) The intra-Layer edges connecting SSs within Layers 0-3. The latter causes the PV-to-PC path taken by some vCompounds to be much longer than the shortest PV-to-PC path illustrated in Fig 3B.

### vCompound-4 does not achieve the cross-system validation target

vC4 represents a highly permeable xenobiotic having an intermediate hepatic extraction ratio. It uses *pBind* = 0.1 and *pMetabolize* = 0.015. The extra-vHPC behaviors of vC3 and vC4 Metabolites are identical. The results in Fig 9A and 9B show that, within experimental variances, for *pEnter* ≥ 0.1, mean peak Entry rates for vCulture (Fig 9C) overpredict corresponding vHuman values, and the magnitude of the overprediction decreases with decreasing values of *pEnter*. As examples, for *pEnter* = 1.0 (0.1), mean peak Entry rates for vCulture (Fig 9C) overpredict corresponding vHuman values by 2.0-fold (1.2-fold). The explanation for those overpredictions highlights systemic differences in the temporal mixing of disposition and intra-vHPC events between the two systems. A fraction of the Dose in Media Space is transferred randomly to 114 Media-Cell Interfaces during comparable early intervals, whereas, for Body, the fraction is transferred randomly to only 45 SS. During vCulture experiments, a vC4 may enter and exit only two spaces, Media-Cell Interface space and Hepatocytes Space, whereas in vLivers, between entering and exiting an SS, a vC4 may enter and exit additional spaces: Core, Interface, Endothelial Cell Space, and Space of Disse. Also, during the average interval that a vC4 requires to enter and exit a vLiver SS, a vC4 may cycle two or more times between Media and Media-Cell Interface space.

**Fig 9.**
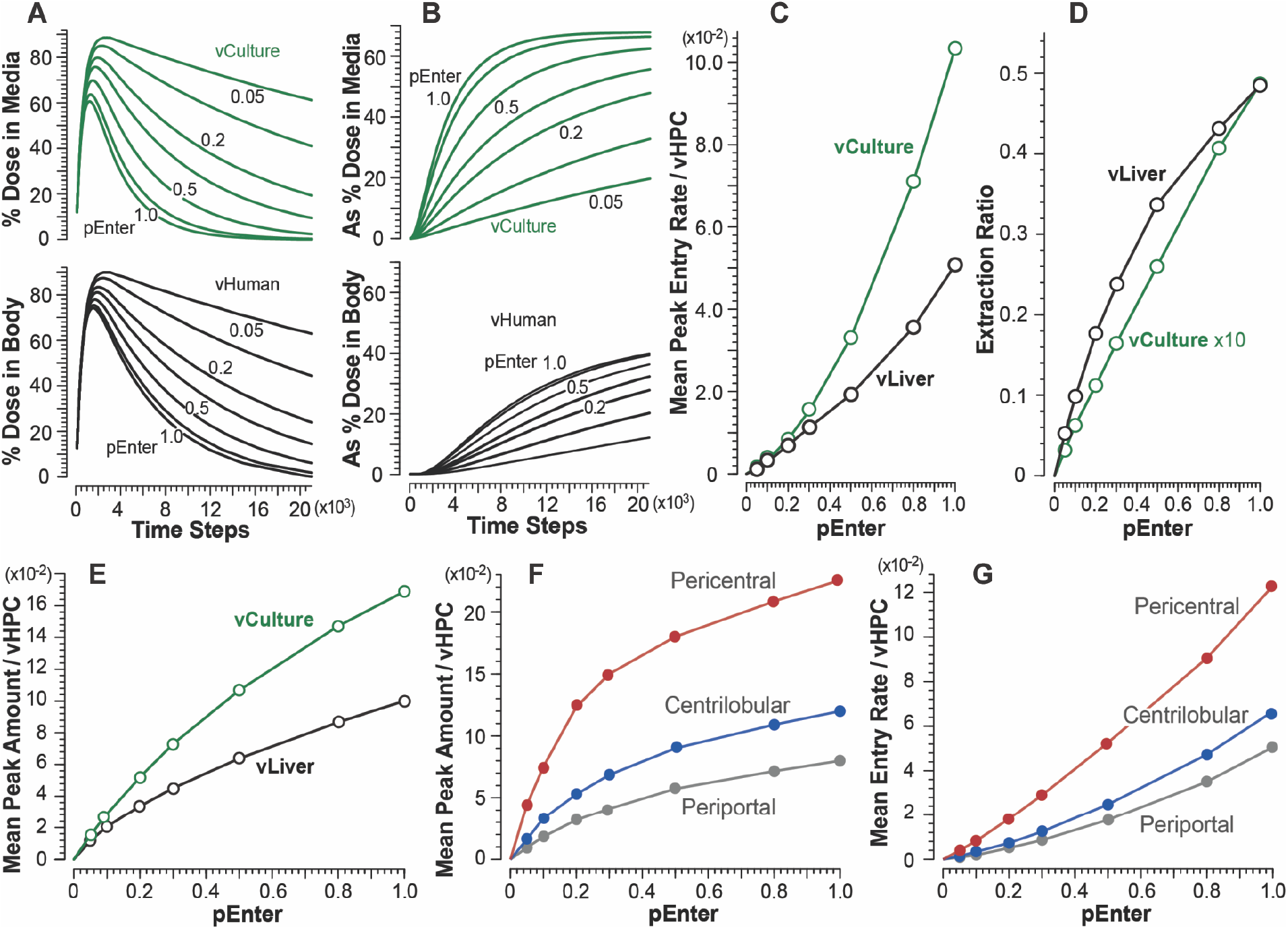
Results from vCulture and vHuman experiments using vC4. (A) Temporal measures of percent Dose in Body (top) and Media (bottom) for each *pEnter.* (B) Amount of Metabolite in Body (top) and Media (bottom) as percent of Dose for each *pEnter.* (C) Correlations between mean peak Entry rates and *pEnter*. (D) Correlations between mean plateau values of Extraction Ratio and *pEnter*. (E) Correlations between systemwide mean peak amounts of vC4 per vHPCs and *pEnter*. (F) Correlations between mean peak amounts of vC4 per vHPCs within the three vLiver bands and *pEnter*. (G) Correlations between mean peak Entry rates within the three bands and *pEnter*.

Entry events and Enzyme binding events are correlated. Results in Fig 9E show that, for vCulture, system-wide mean peak amounts of vC4 per vHPC are larger than corresponding system-wide mean peak amounts in vLiver. Thus, system-wide by corresponding times, more vC4 Metabolism has occurred in vCulture experiments. During comparable intervals, Metabolic events per vHPC within vCulture and within vLiver’s PP band are similar. However, within vLiver, Metabolic events per vHPC increase PP-to-CL-to-CV (Fig 9F and 9G).

In both systems, mean values of Extraction Ratio plateau by about 7,500 TS. The large difference in Extraction Ratios is a consequence of vLiver-vCulture structural differences. In vCulture, a vC4 that is bound to an Enzyme and released later without being Metabolized is, on average, unlikely to become Enzyme-bound and Metabolized within another vHPC before returning to Media. However, in vLiver, a vC4 that is bound to an Enzyme in an upstream vHPC and later released unchanged will, on average, have additional opportunities to be Metabolized before returning to Body, as evidenced by peak amounts per vHPC (Fig 9F) and Entry rates (Fig 9G) increasing from PP to PC bands.

On average, during a vCulture experiment using *pEnter* = 1.0, the mean peak Entry rate is 0.102, whereas during a vHuman experiment, the corresponding values within the PP, CL, and PC bands are 0.051, 0.066, and 0.12, respectively. That PP-to-PC increase is larger for smaller *pEnter* values, similar to the increases measured for vC1 (Fig 5F). To illustrate for *pEnter* = 1.0 (0.1), mean peak Entry rates increase 2.35-fold (3.6-fold) from the PP to the PC band. The patterns for the correlation between mean peak Entry rates within the PP, CL, and PC bands (Fig 9G) are similar to corresponding patterns for vC1 (Fig 5F), although Entry rates of the former are smaller due to upstream binding of vC4s to Enzymes and cumulative Metabolism.

## Discussion

The work presented marks the first step in demonstrating the feasibility of the plan illustrated in Fig 1. Essential for leveraging the potential of Protocol 3 in Fig. 1 is achieving the following vCulture-vHuman cross-system validation target: for specific types of vCompounds, a 1:1 equivalency exists between measures of unbound vCompound Entry rates (per vHPC) made during vCulture and vHuman experiments. Hence, in those cases, vCulture Entry rates directly predict the corresponding measures made during vHuman experiments. For the four vCompound types studied, we achieved the 1:1 equivalency in 15 of 28 cases: 1) for the seven vC1 cases (*pEnter* = 0.05-1.0) where the vCompound removal rate is zero; 2) for the seven vC3 cases where vCompound removal rates are much smaller than vHPC Entry rates (maps to a slowly metabolized xenobiotic); and 3) in one vC2 case where *pEnter* (extra-Cellular unbound fraction) = 1.0 and thus vCompound and removal rates ≅ Entry rates. For the other 13 cases, differences in the structural organization of vHPCs within the vCulture and vHuman systems cause vCulture measures to either underpredict (vC2, *pEnter* ≤ 0.8) or overpredict (vC4, all seven *pEnter* values) corresponding vHuman measures. The magnitude of the vC2 underpredictions increase as *pEnter* decreases (Fig 7C). Using *pEnter* = 0.05 caused a 4.4-fold underprediction.

We recorded and measured temporal differences in model mechanism details as they unfolded within each system during executions. We then used that information to develop explanations for how and why vCulture measures either under- or overpredicted corresponding vHuman measures, which is the purpose of Protocol 3. Those explanations demonstrate how Protocol 3 can facilitate deep thinking about the actual mechanisms responsible for IVIVE discrepancies.

Previous reports detail strengths and weaknesses of the analogical approach and methods employed herein [7,9,10,14,17,28]. We argue that, because of measurement limitations and the fog of multi-source uncertainties impacting IVIVEs, increased reliance on analogical arguments may be necessary to make progress disentangling mechanisms contributing to IVIVE inaccuracies. Bartha provides guidelines for assessing the scientific acceptability and limitations of analogical arguments [18], and reliance on analogical arguments and reasoning can be both a limitation and strength [19].

Most IVIVE methods employ either the well-stirred (most common) or parallel tube liver model even though researchers understand that neither model can represent liver physiology complexities, such as the heterogeneity in enzymatic expression [29,30]. Those liver models continue to be used because doing so limits the mathematical complexities of the methods employed. Given the results of vC1-vC4 experiments, the following conjecture is reasonable. IVIVE methods that abstract away critical structural information about hepatocyte organization within the liver contribute to discrepant IVIVE predictions of hepatic clearance in humans. The complicacy of the vLiver is, in part, the product of an overarching requirement of the plan illustrated in Fig 1; the vHuman and vLiver are designed to be reused, without significant structural change, to achieve Protocol 2 validation targets for a wide variety of referent xenobiotics.

The highly permeable vC1-vC4 are a tiny sample from the space of vCompound types and their xenobiotic referents. Additional work will be needed to explore the relative vCulture-vHuman differences in disposition, removal, and Entry rate measures for a diverse variety of vCompound types. The vCompound disposition parameters, which are specific to a vCompound type, are *pEnter* and *pExit* (each value can be Cell-type-specific), and *forward-* and *lateralBias* (which can vary within different SS Spaces). Within vHPCs, the Enzyme type, *bindersPerCell(Min/Max)*, and the parameters, *pBind, bindCycles, pMetabolize,* and *bindersPerCell(Min/Max)* can be vCompound-type-specific. Transport parameters can be included as part of future explorations and made vCompound-type-specific without requiring vLiver structural changes. vCompound types utilizing different combinations of those parameter values may amplify or diminish the over- and underpredictions described above.

There is ample evidence that in vitro-in vivo differences in hepatocyte heterogeneity contribute significantly to IVIVE prediction discrepancies [3,4,30,31], yet the vCulture and vHuman experiments employed only homogeneous vHPCs. This work provides the necessary and essential foundation for future research that is needed to explore in vitro-in vivo differences in vCompound Entry and removal rates caused by vHPC heterogeneity. Expression levels increase PP-to-PC for most liver genes responsible for detoxification and xenobiotic metabolism [32]. We anticipate that exploration of PP-to-PC difference in vHPC-vCompound interaction properties will be required to meet many xenobiotic-specific Protocol 2 validation targets. The vLiver’s design enables simulating such location-dependent model mechanisms. Smith et al. and Kennedy et al. provide examples [12,23].

Because of the structural organization of vHPCs within vLivers, vHPC Entry rates increase PP-to-PC for all four vCompounds, notwithstanding that all vHPCs are identical. That is true even for vC2, where each Entry event is also a removal event. Additional work is needed to explore whether or not the degree of interaction between Metabolism and Entry rates increasing PP-to-PC will amplify discrepant IVIVE predictions of hepatic clearance.

We anticipate that achieving Protocol 2 validation targets, for many xenobiotics, will require that Metabolism increase PP-to-PC. The resulting increases in model mechanism complexity will require clear evidence that we have not compromised cross-system validation requirements. That evidence will be needed to support the model mechanism-based explanations of the discrepant IVIVE prediction developed during Protocol 3. Inclusion of one (or more) of the vCompounds studied here can provide that evidence. An explanation follows.

In Methods, we note that each Dose includes equal amounts of Marker and vCompound. Marker has served as a multi-attribute internal standard. Note that if we repeat vCulture and vHuman experiments in which Dose = vC2 + Marker but replace Marker with vC1, measures of the temporal dynamics of vC2 in both experiments will be the same (within experiment variance). Further, the corresponding vC1 measures will be the same as those in an experiment dosed with vC1 + Marker. To facilitate achieving Fig 1 use cases, we can replace Marker with a vCompound from the vC1-vC4 set. To illustrate, consider a set of Protocol 2 experiments. Dose contains a 50:50 mix of a new vCompound and vC1 objects (using *pEnter* = 1) in place of Marker. Parameterizations of the new vCompound will be refined iteratively (multiple rounds of systematic parameter adjustments) so that its measured properties become analogous to selected properties of the referent xenobiotic. During that process, vC1 properties are expected to be invariant. Upon concluding each experiment, vC1 measurements, such as those in Fig 4 and Fig 5, are plotted along with new vCompound measurements. Code changes are often necessary during iterative refinement processes (e.g., see [25]). Any significant change in vC1 measurements from one refinement cycle to the next serves as a red flag. In such a case, the problem is corrected, the necessary system verification tests are concluded, and iterative refinement continues. Because Metabolism will be a focus for all Protocol 2 experiments, we anticipate that it will be more informative to use a Metabolized vCompound as the internal standard, such as vC4, rather than vC1. During parameter refinement, the PP-to-PC pattern of Metabolism of a new vCompound can be adjusted, as required, while keeping that pattern invariant for vC4.

Once Protocol 1 and 2 validation targets have been achieved for several xenobiotics, quantitative mappings can be established between a subset of xenobiotic physicochemical property values (or molecular descriptors calculated from structure information) and the set of vCompound-specific parameter values listed above. Using an earlier version of the vLiver, Yan et al. identified different sets of vCompound-specific parameter values that enabled quantitative validation against liver perfusion data for four drugs [5]. The authors then used those relationships to successfully predict vCompound-specific parameter values for two additional drugs, given only values of each drug’s physicochemical properties. The authors discuss the strengths, weaknesses, and limitations of using methods to predict vCompound-specific parameter values, given only values of a new xenobiotic’s physicochemical properties.

In summary, taken together, the results herein support the feasibility of using vCulture and vHuman model mechanisms to help explain observed IVIVE discrepancies. However, considerable additional work is needed to demonstrate feasibility. Near-term, additional work is needed to explore a larger variety of vCompound-specific parameter values and to identify and explain vCulture-vHuman differences in vCompound disposition and Metabolism caused by vHPC heterogeneity.

## Acknowledgements

We thank the authors of the MASON simulation toolkit and also the creators of the SimTK project-hosting platform for facilitating development and distribution of this project.

## Author Contributions

**Conceptualization** CAH, AKS, GEPR

**Data Curation** AKS, GEPR, PK

**Investigation** PK, AKS, LD, CAH

**Methodology** AKS, GEPR, CAH

**Project Administration** AKS, CAH, PK

**Resources** GEPR

**Software** GEPR, AKS

**Supervision** CAH

**Validation** AKS, PK, GEPR

**Visualization** CAH, AKS, PK

**Writing**

**Original Draft Preparation** PK, LD, AKS, CAH, LD, RCK

**Review & Editing** CAH, AKS, PK, LD, RCK, GEPR

## Supporting information captions

**S1 Table. Key vCompound parameter values and specifications for MC experiments and structural features.** Table subsections for vHuman and vCulture: Experiment; vCompound features, events, & activities; vCompound convection and dispersion; and Structural features.

---

**Supporting S1 Table.** Key vCompound parameter values and specifications for MC experiments and structural features. Table subsections for vHuman and vCulture: Experiment; vCompound features, events, & activities; vCompound convection and dispersion; and Structural features.

**Key vCompound parameter values and specifications for MC experiments and structural features.**
| Experiment | vHuman | vCulture |
| --- | --- | --- |
| Number of MC executions | 12 | 12 |
| Number of time steps | 21600 | 21600 |

| vCompound Features, Events, & Activities | vHuman | vCulture |
| --- | --- | --- |
| Marker (Membrane crossing) | FALSE | FALSE |
| vC1 (Membrane crossing) | TRUE | TRUE |
| pEnter | [0.05, 0.1, 0.2, 0.3, 0.5, 0.8, 1.0] | [0.05, 0.1, 0.2, 0.3, 0.5, 0.8, 1.0] |
| pExit | 1 | 1 |
| vC2 (Membrane crossing) | TRUE | TRUE |
| pEnter | [0.05, 0.1, 0.2, 0.3, 0.5, 0.8, 1.0] | [0.05, 0.1, 0.2, 0.3, 0.5, 0.8, 1.0] |
| pExit | 1 | 1 |
| vC3 (Membrane crossing) | TRUE | TRUE |
| pEnter | [0.05, 0.1, 0.2, 0.3, 0.5, 0.8, 1.0] | [0.05, 0.1, 0.2, 0.3, 0.5, 0.8, 1.0] |
| pExit | 1 | 1 |
| vC4 (Membrane crossing) | TRUE | TRUE |
| pEnter | [0.05, 0.1, 0.2, 0.3, 0.5, 0.8, 1.0] | [0.05, 0.1, 0.2, 0.3, 0.5, 0.8, 1.0] |
| pExit | 1 | 1 |

| vCompound Convection, Dispersion | vHuman | vCulture |
| --- | --- | --- |
| innerForwardBias | 0.2 | 0.2 |
| innerLateralBias | 0.5 | 0.5 |
| outerForwardBias | 0.2 | 0.2 |
| outerLateralBias | 0.5 | 0.5 |
| ssFlowRate | 1 | 1 |

| Structural Features | vHuman | vCulture |
| --- | --- | --- |
| Edges | vLiver, Body, and Intro space (to contain Dose)<br>55 Layer 0-to-1; 65 L1-to-L2; 35 L2-to-L3; 25 L3-to-L4 edges.<br>Intra-Layer edges:<br>Layer 0 =20, Layer 1=7, Layer 2=5, Layer 3=2, Layer 4=0 | Not used: Core, Endothelial Cell Space, and Space of Disse<br>Media-Cell Interface = merged (Interface Space, PV, CV) |
| Width (grid spaces) | SS = 15 | Hepatocyte Space = 15 |
| Length (grid spaces) | SS = 5 ± 1(SD) | Hepatocyte Space = 5 |
| MC Sampled per experiment | 12 vLobules | 114 Hepatocyte Spaces |
| vHPC's per execution | Mean = 8,475 (SD = 167) | 8,550 |
| Dose | Single | Single |
| Marker | 50,000 | 50,000 |
| vCompound | 50,000 | 50,000 |
| SVN Version ID | 1352 | 1352 |

